# Antibody evasion and receptor binding of SARS-CoV-2 variants PQ.16.1.1 and RK.1

**DOI:** 10.64898/2026.07.21.739818

**Authors:** Peizhuo He, Yunwen Song, Caiwan Guo, Lingling Yu, Yuanling Yu, Fanchong Jian, Fei Shao, Yunlong Cao

**Affiliations:** Changping Laboratory, Beijing, P. R. China; Chinese Academy of Medical Science & Peking Union Medical College, Beijing, P.R. China; Academy for Advanced Interdisciplinary Studies, Peking University, Beijing, P. R. China; Biomedical Pioneering Innovation Center (BIOPIC), School of Life Sciences, Peking University, Beijing, P. R. China; Peking–Tsinghua Center for Life Sciences, Peking University, Beijing, P.R. China

## Abstract

The recent global expansion of the SARS-CoV-2 variant NB.1.8.1 has driven the emergence of sublineages PQ.16.1.1 and RK.1, which independently acquired the D420N mutation in their receptor-binding domains and and now dominate the Asia-Pacific region. Evaluations utilizing surface plasmon resonance and pseudovirus assays demonstrate that these sublineages exhibit significantly reduced human ACE2 receptor engagement compared to their parental strain. However, this functional cost is offset by a marked ability to evade humoral immunity, specifically demonstrating profound resistance to Class 1 neutralizing monoclonal antibodies and convalescent plasma from Wuhan-Hu-1-primed individuals. This convergent evolution exemplifies a classical viral trade-off, sacrificing receptor binding efficiency to escape population-level immune pressure. Consequently, these findings suggest these variants will soon spread globally and emphasize the critical need for ongoing surveillance to monitor D420N-carrying lineages.

## Results

Since its rapid global spread beginning in late 2024, the SARS-CoV-2 variant NB.1.8.1 has progressively displaced older Omicron variants, establishing near-total dominance in Asia ^1-3^. More recently, two NB.1.8.1-derived sublineages, PQ.16.1.1 and RK.1, have emerged and expanded significantly, particularly in China and Singapore. Specifically, the sublineage PQ.16.1.1 acquired the amino acid substitutions D253G (within the N-Terminal Domain), alongside N417T, D420N, and I478T within the receptor-binding domain (RBD), relative to the parental NB.1.8.1 strain (Figure 1A). Concurrently, the RK.1 sublineage (formally classified as a descendant of the PQ.17.7.2.1 branch) acquired D420N, H445P, and I478T (Table S1). Furthermore, PQ.16.1.1 has continued to evolve into the SV series sublineages (predominantly SV.2 and SV.2.1), which maintained all RBD mutation, including D420N, and have subsequently come to dominate the circulating SARS-CoV-2 strains in Singapore (Figure 1A).

**Figure 1.**
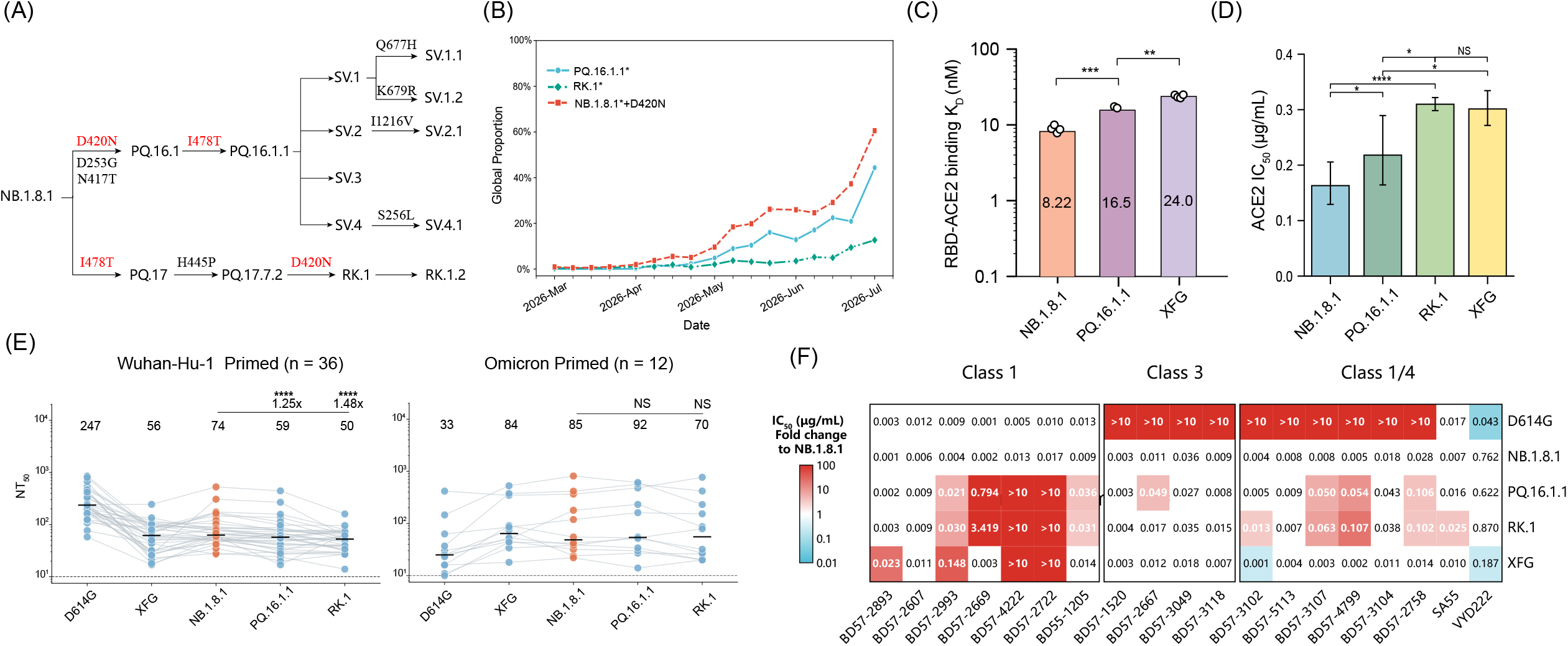
Antigenic and virological characteristics of emerging SARS-CoV-2 variants PQ.16.1.1 and RK.1. (A) Genetic changes in the spike glycoprotein of prevalent SARS-CoV-2 variants; key mutations observed in these variants are marked in red. (B) Global weekly prevalence ratio of PQ.16.1.1^*^, RK.1^*^, NB.1.8.1^*^+ Spike_D420N and others sublineages. Data were collected from Global Initiative on Sharing All Influenza Data database (March, 2026 - July, 2026). (C) The binding affinity of NB.1.8.1, PQ.16.1.1 and XFG RBD proteins to human ACE2, established by SPR. Each circle indicates a technical replicate. Geometric mean K_D_ values (nM) are displayed. Log-transformed data were analysed using a two-tailed t test to compare the means between the two groups. (D) IC_50_ values for the neutralisation of NB.1.8.1, PQ.16.1.1, RK.1 and XFG pseudoviruses by soluble human ACE2. Technical replicates are shown as individual circles. IC_50_(μg/mL) was log-transformed before two-tailed t test analysis. (E) NT_50_ values were determined using convalescent plasma samples collected from Wuhan-Hu-1 primed individuals (n = 36) and Omicron-primed individuals (n = 12). Cohort labels and sample sizes are provided above each panel. The dashed line represents the detection limit (NT_50_=10). Geometric means, fold reductions, and corresponding p values (Wilcoxon rank-sum test) are noted above each group. (F) IC_50_ values (μg/mL) of monoclonal neutralising antibodies targeting RBD epitopes against NB.1.8.1, PQ.16.1.1, RK.1 and XFG variants. Fold changes relative to NB.1.8.1 are shown by background colour: red denotes increased resistance and blue denotes enhanced sensitivity. Colour scale indicates the fold-change magnitude. IC_50_=50% inhibitory concentration. NS=not significant. NT_50_=50% neutralisation titre. RBD=receptor-binding domain. Wuhan-Hu-1 reference strain. *p<0.05. **p<0.01. ***p<0.001. ****p<0.0001.

The independent, convergent acquisition of D420N, a mutation that previously predicted to cause immune evasion of Class 1 neutralizing antibodies ^4-8^, across distinct evolutionary branches of the NB.1.8.1 family tree strongly indicates powerful, population-level positive selection. Global genomic surveillance data collected between March and July 2026, utilizing sequences submitted to the Global Initiative on Sharing All Influenza Data (GISAID) and the National Center for Biotechnology Information (NCBI) virus databases, demonstrate that D420N-carrying NB.1.8.1 sublineages have successfully achieved regional dominance across the Asia-Pacific and are progressively displacing other circulating variants (Figure 1B). Here, we investigated the receptor engagement and humoral evasion characteristics of these D420N mutation carrying variants.

We first assessed the human angiotensin-converting enzyme 2 (hACE2) receptor engagement of these variants. Surface plasmon resonance (SPR) demonstrated that the recombinant PQ.16.1.1 RBD exhibited a significantly reduced binding affinity for hACE2 compared to NB.1.8.1, as reflected by an increased equilibrium dissociation constant (K_D_) (Figure 1C, Figure S1 and Table S2). Notably, the RBD–hACE2 binding affinity of PQ.16.1.1 remains similar to or stronger than that of XFG. Given that XFG has been circulating effectively, this indicates that the D420N mutation is not detrimental to ACE2 receptor binding. Next, we generated vesicular stomatitis virus pseudotypes bearing the spike proteins of these variants and measured their susceptibility to soluble hACE2-mediated inhibition, an assay that measures the receptor-binding strength and engagement efficiency of the variant’s Spike. Consistently, both PQ.16.1.1 and RK.1 showed significantly reduced spike-mediated ACE2 engagement relative to NB.1.8.1; however, their engagement efficiency remained similar to or stronger than that of XFG (Figure 1D). Collectively, these results suggest that while D420N reduces receptor engagement efficiency, it remains within the range required for effective viral prevalence.

Next, we evaluated humoral immune evasion using human convalescent plasma and a panel of RBD-targeting neutralizing monoclonal antibodies (mAbs). The individual plasmas were collected between April and August 2025 in Beijing, China, when NB.1.8.1 was dominant (Table S3). Compared with NB.1.8.1, both PQ.16.1.1 and RK.1 exhibited significantly reduced neutralization titers against Wuhan-Hu-1-primed plasma (from individuals who received Wuhan-Hu-1 inactivated vaccines). In contrast, susceptibility to Omicron-primed plasma (from unvaccinated individuals whose first exposure was to Omicron) remained insignificant. This highlights that the D420N mutation may specifically exploit vulnerabilities in the antibody response induced by Wuhan-Hu-1 priming (Figure 1E).

Indeed, mAb profiling revealed that both PQ.16.1.1 and RK.1 substantially resisted Class 1 antibodies, with RK.1 demonstrating a more extensive loss of neutralization (Figure 1F). This aligns with predictive models identifying residue 420 as an immune-pressure hotspot within the Class 1 epitope, which physically overlaps the ACE2-binding footprint ^4-8^. Combined with the reduced ACE2 engagement efficiency of RK.1 compared to PQ.16.1.1, this also indicates that 417N (present in RK.1) exhibits stronger Class 1 antibody evasion compared to 417T (present in PQ.16.1.1) at the cost of ACE2 binding strength, given that residue 417 is their primary mutational difference on the RBD.

Compared to Class 1 mAbs, PQ.16.1.1 and RK.1 demonstrated weak to no antibody evasion against Class 3 and Class 1/4 mAbs. The elicitation of these Omicron-specific antibody classes varies sharply across populations and dominates the antibody response in Omicron-primed individuals ^6-8^. This observation aligns with the fact that PQ.16.1.1 and RK.1 did not cause as substantial immune evasion in Omicron-primed cohorts compared to Wuhan-Hu-1-primed cohorts (Figure 1E).

In summary, the convergent acquisition of the D420N substitution in NB.1.8.1 sublineages again illustrates a classic SARS-CoV-2 RBD evolution trade-off: a sacrifice in hACE2 receptor engagement in exchange for profound, targeted evasion of Class 1 neutralizing antibodies. This phenomenon has been frequently observed in the evolutionary trends of Omicron, as represented recently by the evolution of JN.1 sublineages ^9-10^. Interestingly, the 420N mutation did not occurr in the XFG backbone. This is because XFG’s ACE2 binding is far weaker than that of NB.1.8.1, making XFG lacks the buffering room necessary for the evolution of 420N. Given the increased evasion of Class 1 antibodies by these D420N-carrying sublineages, it is highly likely that these variants will spread from Asia and begin to prevail in Western countries, as mRNA-vaccinated individuals are enriched with Class 1 neutralizing antibodies. This underscores the need for continued, stringent global surveillance of those SARS-CoV-2 variants, especially for future compensatory mutations that could restore receptor engagement while preserving this newly acquired immune evasion mutation.

## Acknowledgments

We extend our gratitude to the scientific community for their continued efforts in monitoring SARS-CoV-2 variants, as well as to all volunteers who contributed blood samples for this study. This project is financially supported by Changping Laboratory (2026D-04-01 to Y.C.).

## Author Contributions

Y.C. designed and supervised the study. P.H. and Y.C. wrote the manuscript with input from all authors. P.H., F.J., and C.G. performed the sequence analyses and prepared the illustrations. Y.Y. constructed the pseudoviruses. L.Y., Y.S., and F.S. processed the plasma samples and performed the pseudovirus neutralisation assays. P.H. and Y.C. analysed the neutralisation data.

## Methods Details

### Global lineage prevalence analysis

SARS-CoV-2 sequence metadata were retrieved from the GISAID EpiCoV database on July 2026^1^. All available records with collection dates between March and July 2026 were included in the initial dataset. Pango lineage assignments provided in the GISAID metadata were used for preliminary classification. To account for lineage aliases and newly designated descendants, lineage_notes.txt and alias_key.json were obtained from the cov-lineages/pango-designation GitHub repository and used to resolve aliased lineage names and compile the complete sets of descendants of PQ.16.1.1, RK.1, and NB.1.8.1. An asterisk denotes a parental lineage and all of its descendants. NB.1.8.1* sequences carrying the Spike_D420N substitution were identified from the amino-acid substitution annotations in the GISAID metadata. Records lacking a complete collection date or an informative Pango lineage assignment were excluded. Sequences were grouped by week according to their collection dates and assigned to categories in the following order: PQ.16.1.1*, RK.1* and NB.1.8.1* sequences carrying Spike_D420N. For each week, the global proportion of a category was calculated as the number of sequences assigned to that category divided by the total number of eligible sequences collected during the same week and was expressed as a percentage.

### Surface Plasmon Resonance

Surface plasmon resonance (SPR) measurements were performed using a Biacore 8K system (12914229; Cytiva). Human ACE2 protein was immobilized on Protein A sensor chips (29127556; Cytiva). Purified receptor-binding domains (RBDs) from SARS-CoV-2 variants were serially diluted to 6.25, 12.5, 25, 50, and 100 nM and subsequently injected over the sensor-chip surface. Binding responses were recorded at room temperature using Biacore 8K Control Software (version 4.0.8.19879; Cytiva). The resulting sensorgrams were fitted to a 1:1 binding model, and equilibrium dissociation constants (K_D_) were calculated using Biacore 8K Evaluation Software (version 4.0.8.20368; Cytiva).

### Patient recruitment and plasma isolation

Participants in the Random cohort donated peripheral blood after providing written informed consent. Consent covered the collection and storage of blood specimens, their use for research purposes, and publication of data arising from the study. The protocol was reviewed and approved by the Tianjin Municipal Health Commission and the Ethics Committee of Tianjin First Central Hospital (Ethics Committee Archiving No. KEYAN20241022-2) and by the Ethics Committee of Beijing Ditan Hospital, Capital Medical University (Ethics Committee Archiving No. DTEC-KY2024-112-01). The study followed the principles set out in the Declaration of Helsinki.

For sample preparation, whole blood was diluted with an equal volume of PBS containing 2% fetal bovine serum. Ficoll density-gradient centrifugation (Cytiva, Cat. No. 17-1440-03) was then used to separate the plasma and PBMC fractions. Plasma was recovered from the upper layer, aliquoted, and maintained at -20°C or colder until analysis. Before experimental testing, all plasma aliquots were heat-inactivated.

### Pseudovirus neutralisation assay

SARS-CoV-2 variant spike pseudoviruses were generated using a vesicular stomatitis virus (VSV)-based packaging system. Briefly, 293T cells (American Type Culture Collection [ATCC], CRL-3216) were transfected with the corresponding spike-expression plasmids together with G*ΔG-VSV (VSV-G-pseudotyped virus; Kerafast). Following culture, pseudovirus-containing supernatants were harvested, filtered, divided into aliquots, and stored at −80 °C until use.

For the neutralization assay, the prepared pseudoviruses were incubated with serially diluted monoclonal anti-S-RBD antibodies, human ACE2-Fc, or plasma samples in 96-well plates at 37°C under 5% CO_2_ for 1 h. Plasma samples were initially diluted 1:10 and then subjected to five consecutive Teefold dilutions. Monoclonal antibodies and ACE2-Fc were first adjusted to 0.1 mg/mL; the monoclonal antibodies were then diluted 100-fold, whereas ACE2-Fc was diluted 50-fold. Both were subsequently subjected to five consecutive fivefold dilutions. Dissociated Huh-7 cells (Japanese Collection of Research Bioresources [JCRB], 0403) were subsequently added to each well. After incubation for 24 h, the culture supernatants were removed and D-luciferin reagent (Vazyme, DD1209-03) was added. Plates were kept in the dark for 2 min, after which luminescence was recorded using a microplate spectrophotometer (PerkinElmer, HH3400). Half-maximal inhibitory concentration (IC_50_) values were determined by fitting the neutralization data to a four-parameter logistic regression model.

## Supplementary Tables

**Table S1.**
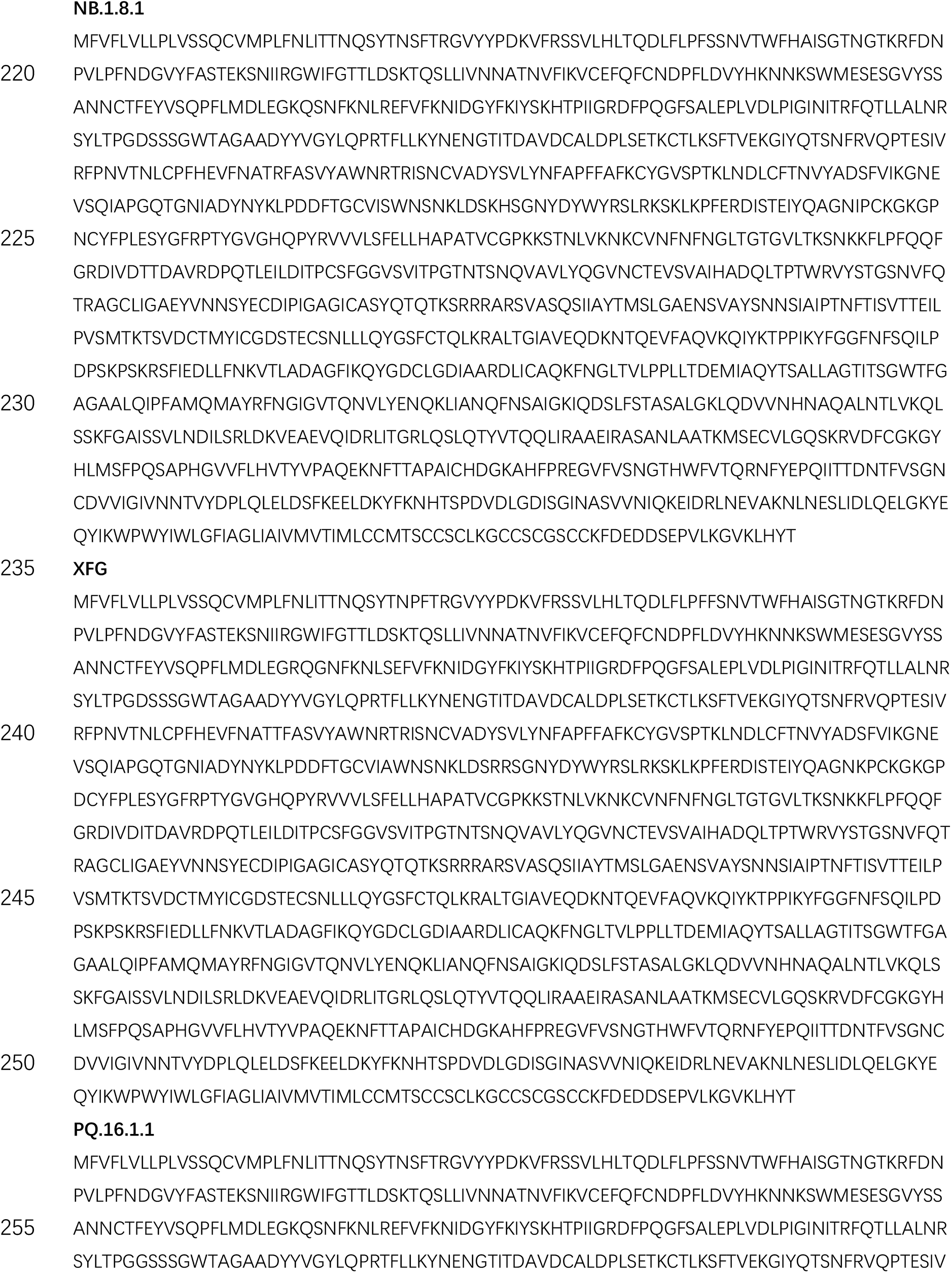
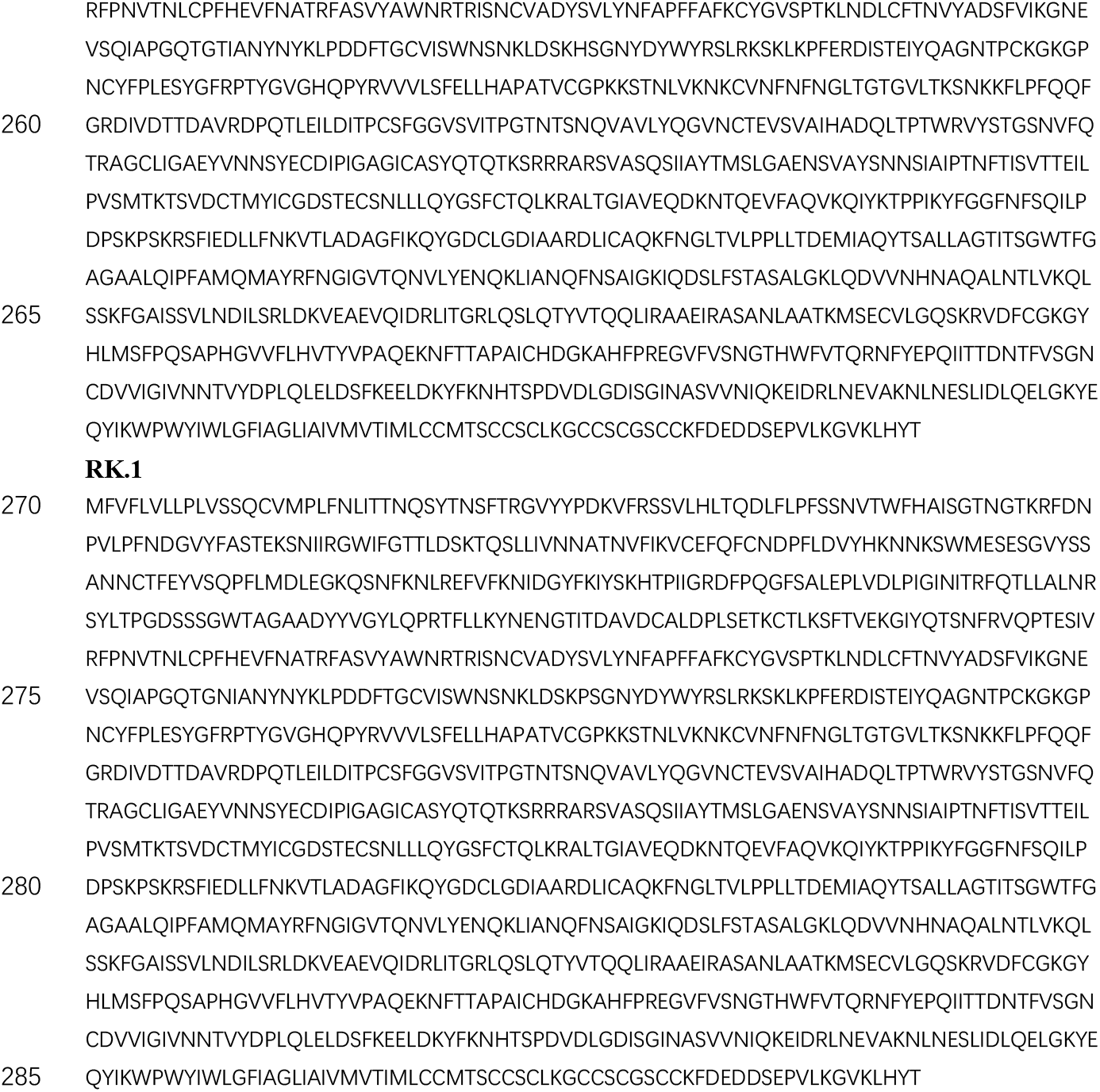
Spike protein sequences of SARS-CoV-2 variants.

**Table S2.** Binding kinetics of SARS-CoV-2 variant RBDs to hACE2 determined by SPR.

| Sample | $k_a$ ($M^{-1} s^{-1}$) | $k_d$ ($s^{-1}$) | $k_d$ ($s^{-1}$) | Rmax (RU) |
| --- | --- | --- | --- | --- |
| NB.1.8.1-RBD | 3.95E+05 | 3.28E-03 | 8.30E-09 | 68.677 |
| NB.1.8.1-RBD | 3.86E+05 | 3.60E-03 | 9.33E-09 | 66.802 |
| NB.1.8.1-RBD | 4.06E+05 | 2.95E-03 | 7.28E-09 | 69.500 |
| NB.1.8.1-RBD | 3.94E+05 | 3.20E-03 | 8.11E-09 | 68.703 |
| PQ.16.1.1-RBD | 9.17E+04 | 1.55E-03 | 1.69E-08 | 61.961 |
| PQ.16.1.1-RBD | 1.65E+05 | 2.65E-03 | 1.61E-08 | 57.213 |
| XFG-RBD | 2.17E+05 | 5.40E-03 | 2.49E-08 | 70.653 |
| XFG-RBD | 1.91E+05 | 4.46E-03 | 2.33E-08 | 73.176 |
| XFG-RBD | 2.40E+05 | 5.45E-03 | 2.27E-08 | 71.490 |
| XFG-RBD | 2.23E+05 | 5.59E-03 | 2.51E-08 | 69.960 |

**Table S3.** Information of SARS-CoV-2 convalescent patients involved in the study.

| Sample Type | Gender | Age | Wuhan-1 primed | Sampling time point | NT<sub>50</sub> |  |  |  |  |
| --- | --- | --- | --- | --- | --- | --- | --- | --- | --- |
|  |  |  |  |  | D614G | NB.1.8.1 | PQ.16.1.1 | RK.1 | XFG |
| Omicron primed individuals | female | 39 | no | 2025/4/9 | 417 | 365 | 480 | 768 | 346 |
| Omicron primed individuals | female | 47 | no | 2025/4/9 | 30 | 179 | 232 | 87 | 50 |
| Omicron primed individuals | male | 75 | no | 2025/5/14 | 25 | 24 | 27 | 26 | 75 |
| Omicron primed individuals | female | 38 | no | 2025/5/28 | 16 | 55 | 53 | 79 | 91 |
| Omicron primed individuals | female | 21 | no | 2025/6/26 | 63 | 119 | 161 | 153 | 85 |
| Omicron primed individuals | female | 42 | no | 2025/7/2 | 37 | 416 | 567 | 418 | 375 |
| Omicron primed individuals | female | 79 | no | 2025/7/16 | 25 | 32 | 30 | 20 | 45 |
| Omicron primed individuals | male | 51 | no | 2025/7/16 | 16 | 22 | 14 | 20 | 18 |
| Omicron primed individuals | female | 52 | no | 2025/7/24 | 23 | 39 | 34 | 20 | 53 |
| Omicron primed individuals | female | 42 | no | 2025/8/13 | 11 | 38 | 55 | 27 | 34 |
| Omicron primed individuals | female | 68 | no | 2025/8/13 | 146 | 813 | 611 | 231 | 529 |
| Omicron primed individuals | female | 43 | no | 2025/8/13 | 10 | 42 | 52 | 32 | 44 |
| Wuhan-Hu-1 primed individual | female | 28 | yes | 2025/4/9 | 854 | 147 | 137 | 66 | 88 |
| Wuhan-Hu-1 primed individual | female | 57 | yes | 2025/4/16 | 199 | 52 | 34 | 29 | 64 |
| Wuhan-Hu-1 primed individual | female | 55 | yes | 2025/4/16 | 320 | 79 | 57 | 79 | 98 |
| Wuhan-Hu-1 primed individual | female | 40 | yes | 2025/4/16 | 390 | 60 | 62 | 32 | 54 |
| Wuhan-Hu-1 primed individual | female | 45 | yes | 2025/4/16 | 500 | 171 | 114 | 54 | 86 |
| Wuhan-Hu-1 primed individual | female | 31 | yes | 2025/4/23 | 77 | 27 | 30 | 29 | 43 |
| Wuhan-Hu-1 primed individual | female | 25 | yes | 2025/4/17 | 192 | 51 | 48 | 57 | 44 |
| Wuhan-Hu-1 primed individual | male | 36 | yes | 2025/4/24 | 239 | 44 | 27 | 30 | 31 |
| Wuhan-Hu-1 primed individual | male | 28 | yes | 2025/4/24 | 58 | 55 | 58 | 32 | 17 |
| Wuhan-Hu-1 primed individual | female | 36 | yes | 2025/4/24 | 242 | 91 | 77 | 79 | 89 |
| Wuhan-Hu-1 primed individual | female | 56 | yes | 2025/4/24 | 533 | 52 | 49 | 59 | 91 |
| Wuhan-Hu-1 primed individual | male | 61 | yes | 2025/5/14 | 426 | 28 | 24 | 14 | 23 |
| Wuhan-Hu-1 primed individual | female | 39 | yes | 2025/5/14 | 242 | 529 | 444 | 160 | 245 |
| Wuhan-Hu-1 primed individual | female | 35 | yes | 2025/5/23 | 250 | 66 | 68 | 47 | 31 |
| Wuhan-Hu-1 primed individual | female | 31 | yes | 2025/5/15 | 174 | 45 | 70 | 41 | 28 |
| Wuhan-Hu-1 primed individual | female | 36 | yes | 2025/5/15 | 161 | 58 | 40 | 41 | 67 |
| Wuhan-Hu-1 primed individual | male | 27 | yes | 2025/5/15 | 180 | 53 | 44 | 38 | 27 |
| Wuhan-Hu-1 primed individual | female | 48 | yes | 2025/5/15 | 250 | 87 | 78 | 61 | 118 |
| Wuhan-Hu-1 primed individual | female | 75 | yes | 2025/5/28 | 672 | 307 | 268 | 62 | 27 |
| Wuhan-Hu-1 primed individual | male | 43 | yes | 2025/5/28 | 207 | 76 | 73 | 74 | 107 |
| Wuhan-Hu-1 primed individual | female | 31 | yes | 2025/6/26 | 175 | 68 | 53 | 50 | 95 |
| Wuhan-Hu-1 primed individual | male | 23 | yes | 2025/6/26 | 108 | 121 | 76 | 66 | 89 |
| Wuhan-Hu-1 primed individual | female | 69 | yes | 2025/7/2 | 158 | 104 | 80 | 73 | 108 |
| Wuhan-Hu-1 primed individual | female | 11 | yes | 2025/7/2 | 391 | 84 | 61 | 48 | 55 |
| Wuhan-Hu-1 primed individual | female | 18 | yes | 2025/7/4 | 467 | 85 | 89 | 52 | 101 |
| Wuhan-Hu-1 primed individual | male | 43 | yes | 2025/7/11 | 464 | 35 | 20 | 27 | 21 |
| Wuhan-Hu-1 primed individual | male | 20 | yes | 2025/7/18 | 210 | 51 | 42 | 40 | 52 |
| Wuhan-Hu-1 primed individual | male | 57 | yes | 2025/7/24 | 158 | 60 | 17 | 65 | 32 |
| Wuhan-Hu-1 primed individual | female | 54 | yes | 2025/8/1 | 333 | 175 | 158 | 80 | 69 |
| Wuhan-Hu-1 primed individual | male | 26 | yes | 2025/8/1 | 201 | 50 | 35 | 40 | 60 |
| Wuhan-Hu-1 primed individual | female | 28 | yes | 2025/8/8 | 122 | 56 | 54 | 75 | 50 |
| Wuhan-Hu-1 primed individual | female | 25 | yes | 2025/8/8 | 165 | 78 | 62 | 72 | 135 |
| Wuhan-Hu-1 primed individual | female | 23 | yes | 2025/8/8 | 421 | 60 | 31 | 62 | 26 |
| Wuhan-Hu-1 primed individual | female | 39 | yes | 2025/8/8 | 801 | 164 | 98 | 96 | 72 |
| Wuhan-Hu-1 primed individual | female | 48 | yes | 2025/8/26 | 234 | 42 | 28 | 27 | 18 |
| Wuhan-Hu-1 primed individual | female | 37 | yes | 2025/8/26 | 125 | 69 | 55 | 34 | 80 |

**Figure S1.**
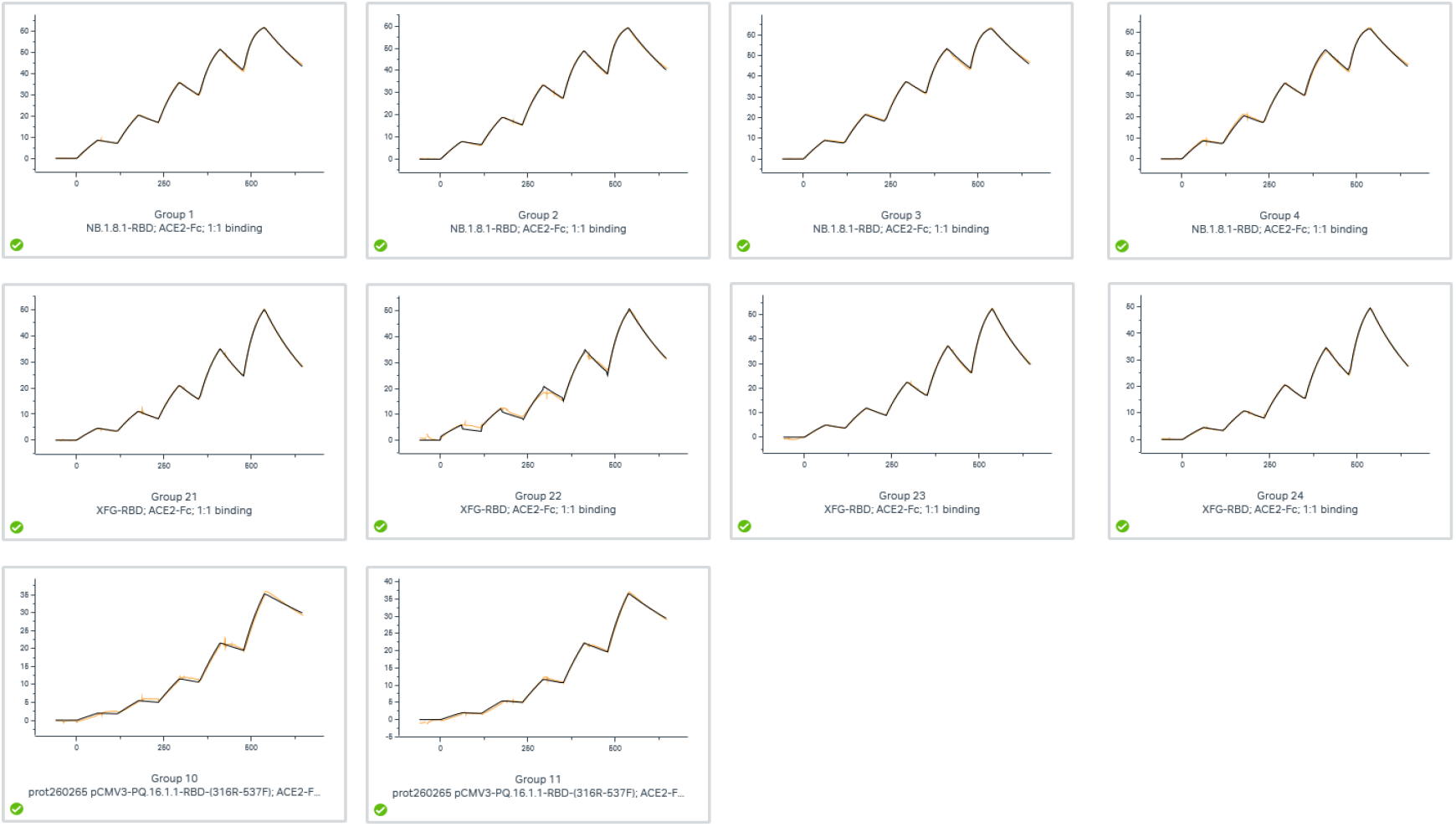
SPR sensorgrams for affinity of hACE2 and SARS-CoV-2 mutants RBD. Representative sensorgrams obtained from single-cycle kinetics measurements performed on a Biacore 8K system. SARS-CoV-2 variant RBDs were serially diluted and sequentially injected over ACE2-immobilized Protein A sensor chips at room temperature. Colored lines represent experimental data, and black lines represent fitted curves based on a 1:1 binding model. Association (ka), dissociation (kd) rate constants, and equilibrium dissociation constant (K_D_) were calculated using Biacore 8K Evaluation Software (version 4.0.8.20368).

## References

1. Guo C, Yu Y, Liu J et al. Antigenic and virological characteristics of SARS-CoV-2 variants BA.3.2, XFG, and NB.1.8.1. The Lancet Infectious Diseases, 2025; 25, e374–e377

2. Mellis IA, Wu M et al. Antibody evasion and receptor binding of SARS-CoV-2 LP.8.1.1, NB.1.8.1, XFG, and related subvariants. Cell Rep. 2025 Oct 28;44(10):116440.

3. Uriu, K., Okumura, K., Uwamino, Y. et al. Virological characteristics of the SARS-CoV-2 NB.1.8.1 variant. The Lancet Infectious Diseases. 2025; 25:e443.

4. Starr, T.N., Greaney, A.J., Hilton, S.K. et al. Deep Mutational Scanning of SARS-CoV-2 Receptor Binding Domain Reveals Constraints on Folding and ACE2 Binding. Cell. 2020; 182:1295–1310.e20.

5. Greaney AJ, Starr TN, Gilchuk P, et al. Complete mapping of mutations to the SARS-CoV-2 spike receptor-binding domain that escape antibody recognition. Cell Host Microbe. 2021;29(1):44–57.e9.

6. Jian, F., Wang, J., Yisimayi, A. et al. Evolving antibody response to SARS-CoV-2 antigenic shift from XBB to JN.1. Nature. 2025; 637:921–929.

7. Cao, Y., Jian, F., Wang, J. et al. Imprinted SARS-CoV-2 humoral immunity induces convergent Omicron RBD evolution. Nature. 2023, 614:521–529.

8. Yisimayi, A., Song, W., Wang, J. et al. Repeated Omicron exposures override ancestral SARS-CoV-2 immune imprinting. Nature. 2024; 625:148–156.

9. Yang, S., Yu, Y., Xu, Y. et al. Fast evolution of SARS-CoV-2 BA.2.86 to JN.1 under heavy immune pressure. The Lancet Infectious Diseases. 2023; 24:e70–e72.

10. Liu, J., Yu, Y., Yang, S. et al. Virological and antigenic characteristics of SARS-CoV-2 variants LF.7.2.1, NP.1, and LP.8.1. The Lancet Infectious Diseases. 2025; 25:e128–e130.

## Supplementary References

1. Shu, Y ., John, M.C. GISAID: Global initiative on sharing all influenza data–from vision to reality. Euro Surveill. 2017; 22: 30494.

